# Varietal differences in physiological and biochemical responses to salinity stress in six finger millet plants

**DOI:** 10.1101/789883

**Authors:** Asunta Mukami, Alex Ng’etich, Easter Syombua, Richard Oduor, Wilton Mbinda

## Abstract

Finger millet is one of the most important cereals that are often grown in semiarid and arid regions of East-Africa. Salinity is known to be a major impediment for the crop growth and production. This study was aimed to understand the mechanisms of physiological and biochemical responses to salinity stress of Kenyan finger millet varieties (GBK043137, GBK043128, GBK043124, GBK043122, GBK043094, GBK043050) grown across different agroecological zones under NaCl-induced salinity stress. Seeds were germinated on the sterile soil and treated using various concentrations of NaCl (100, 200 and 300 mM) for two weeks. Again, the early-seedling stage of germinated plants was irrigated with the same salt concentrations for 60 days. Results indicated depression in germination percentage, shoot and root growth rate, leaf relative water content, chlorophyll content contents, leaf K^+^ concentration, and leaf K^+^/Na^+^ ratios increased salt levels. Contrary, proline and malonaldehyde (MDA) contents reduced sugar content and leaf total proteins. At the same time, the leaf Na^+^ and Cl^−^ amounts of all plants increased substantially with rising stress levels. Clustering analysis revealed that GBK043094 and GBK043137 were placed together and identified as salt-tolerant varieties based on their performance under salt stress. Overall, our findings indicated a significant varietal variability for most of the parameters analysed. These superior varieties identified could be potentially used as promising genetic resources in future breeding programmes development directed towards salt-tolerant finger millet hybrids. Further analysis at genomic level need to be undertaken to better understand the genetic factors that promote salinity tolerance in finger millet.

## Introduction

Salinity is the severest environmental stress that adversely affects the growth of plants and their productivity worldwide (Qadir et al. 2014). This abiotic stress mostly characterizes arid and semiarid regions that experience low rainfall and scarcity of good quality water. Salts accumulation in irrigation without proper drainage water system, coupled with underlying rocks rich with high salts contents, leads to gradual salinization of arable land, thereby affecting soil characteristics. The problem is expected to aggravate further owing to effects of rising sea levels and climate change (Tedeschi et al., 2011). It is estimated that if existing salinity stress phenomenon will continue to persist, more than 50% of the current cultivated agricultural land could be lost by the year 2050 (Wang et al., 2003). As of 2013, the global losses in agricultural production due to lands afflicted by salinity had touched US$12 billion and have been steadily rising ever since (Shabala, 2013).

Over time, plants have evolved complex salt tolerance adaptive mechanisms to counteract the harmful effects of salinity through activation of morphological, physiological biochemical, cellular and molecular responses which include changes to metabolic systems, nutritional disproportion, variation and disorder of membranes and reduction in rate of cell division and growth (Munns et al., 2006; Zhu, 2003). Another critical repercussion of salinity stress to plants is the overproduction of reactive oxygen species (ROS) from the pathways such as photosynthesis, mitochondrial respiration and photorespiration. ROS toxicity comes from their reactions with various cell units, which precipitates an avalanche of oxidative reactions and results to enzymes inactivation, protein degradation, lipid peroxidation and DNA damage (reference). Collectively, these effects inhibit growth and development and subsequently reduce crop yields. The most effective way to combat against salinity is the development of the resistant and tolerant crop varieties. It is therefore paramount to identify the genetic resources with high tolerance, and to understand the mechanisms of salinity tolerance in crops.

The resistance and response of plants to a salt stress varies according to its species or varieties or genotypes or variety and environment which could be attributed to the biological dissimilarities between the species or varieties or genotypes, plant growth, and composition and concentration of the salt stress conditions (Bertazzini et al., 2018; Filichkin et al., 2018; Shabala et al., 2013). Many reports have shown that short term salinity stress significantly affects the germination rate, seedling and root growth as well as ion composition, levels of relative water content, photosynthetic pigments, proline content, level of membrane lipid peroxidation as well as the amounts of reducing sugars and total protein (Dugasa et al., 2019; Sarabi et al., 2017; Kumar and Khare, 2016). These physiological and biochemical indices multivariate cluster analysis been used for classification of salt-sensitive and salt-tolerant varieties so that they can further be used in plant breeding programmes. The prevalent approach to assess the performance of plants against salinity under laboratory conditions is through assessing their physiological and biochemical responses on the application of different concentrations NaCl.

Finger millet, *Eleusine coracana* L. (Geartn), is the 4^th^ most important member of millets after sorghum, pearl millet and foxtail millet, making it one of the most valuable food cereals cultivated in arid and semi-arid regions of Asia and Africa (Chivenge et al., 2015). The crop is well adapted to heat, drought and poor soil stress in marginal and degraded soils. Further, its cereals have comparatively better antioxidant and nutraceutical properties and superb storage qualities which lack in other cereals (Kumar et al., 2016). All these attributes make finger millet one of the important and promising plant genetics resources for agriculture, and food and nutritional security and alleviation of poverty of poor farmers who live in arid, infertile and marginal lands. Despite its importance, finger millet potential yields are adversely affected salinity stress. More specifically, during seed germination and seedling establishment terminal growth phases are extremely susceptible to salinity stress (Ibrahim, 2016; Hema et al., 2014; Zhang et al., 2014). Hence, it is imperative to screen for varieties with intrinsic salinity tolerance for yield improvement breeding programmes. Salinity tolerance during germination and seedling development is crucial for the establishment of plants growing in saline soils of arid and semi-arid regions (Tlig et al., 2008). Accordingly, understanding the physiological and biochemical salinity responses in finger millet is, therefore, of importance in breeding salt resistant and tolerant crops. Owing to the wide disparity in agroecological regions across finger millet growing regions, several finger millet landraces exhibit an adaptation to a large range of environmental conditions and subsequently, represent valuable source of useful genetic source that can be exploited to improve salinity tolerance of finger millet varieties belong to distinct geographical zones in Kenya. We therefore investigated the physiological and biochemical responses to salinity stress of six finger millet varieties under NaCl induced salinity stress.

## Materials and methods

### Plant material, treatments and germination assays

Six Kenyan farmers preferred finger millet varieties (GBK043124, GBK043122, GBK043137, GBK043128, GBK043094 and GBK043050) obtained from Kenya Agricultural and Livestock Research Organization, Gene Bank, Muguga, Kenya were used in this study. Prior to assays, the seeds were sorted by handpicking of the healthy ones, then washed with distilled water to remove dust and other particles. Germination assay was performed using 10 seeds of each variety and at different concentrations of NaCl (100,200 and 300 mM). Seeds were planted in germination trays in round pots containing sterile soil to a depth of approximately 1 cm. The control seeds were irrigated with distilled water. Salinity stress was imposed on treatment groups by irrigating the seeds with various concentrations of NaCl at an interval of 3 days for two weeks. Observations on the rate of germination were scored on the 17^th^ day of treatment.

### Growth conditions under salinity treatment

Germinated finger millet seedlings were grown for 2 weeks under greenhouse conditions of 25±2 °C and 60-70% humidity, with a 16/8-h photoperiod provided by natural sunlight. To assay salinity stress effects on growth of finger millet, the seedlings were subjected to stress by irrigating with NaCl (100, 200 and 300 mM) for 21 days at an interval of 3 days. Control plants were watered with distilled water. In each experiment, five replications were used for each set of treatment. After treatment, five plants from each treatment were sampled at random and the growth of the plants studied by recording the shoot length and root length.

### Relative water content

One leaflet from the first fully expanded leaf of five plants per variety and per treatment was cut from a plant on the 21^st^ day. Immediately after cutting, the leaflet was weighed to obtain the fresh weight (FW). Thereafter, the leaflet was immersed in deionized water under normal room temperature for 4 hours. Afterwards, the leaflet was taken out, thoroughly wiped to remove the water on the blade surface and its weight measured to obtain turgid weight (TW). the leaflet was afterwards dried in an oven for 24 hours and its dry weight (DW) measured. The relative water content (RWC %) was calculated using the formula: RWC = [(FW -DW)/ (TW - DW)] x 100.

### Determination of chlorophyll content

Chlorophyll a, b and total chlorophylls (a + b) were determined according to Arnon (1949). 0.2g of fresh leaves were taken from 21 days-old NaCl (0-300mM) treated plants, finely ground by vortexing several times to remove chlorophyll efficiently. The extract was centrifuged at 5000 g for 3 minutes. The absorbance of the obtained supernatants was measured at 645 and 663 nm using 1240 UV-Vis Spectrophotometer (Shimadzu, Kyoto, Japan). The total chlorophyll content in each sample, expressed in mg/g fresh mass (FM) was calculated using Arnon’s 1949 formula: TC=20.2(A645) □ 8.02(A663) ×V/1000×W where V corresponds to the volume of total extract per litre and W is the mass of the fresh material.

### Proline content measurement

Proline accumulation was determined as described by Bates et al. (1973). Fifty milligrams of fresh leaf tissues from each variety and treatment was homogenized in 10 ml of 3% w/v sulphosalicylic acid and the homogenate was filtrated. The resulting solution was mixed solution of acidic ninhydrin [40% (w/v) acidic ninhydrin (8.8 µM ninhydrin, 10.5 M glacial acetic acid, 2.4 M orthophosphoric acid), 40% (v/v) glacial acetic acid and 20% (v/v) of 3%(v/v) sulphosalicylic acid]. Thereafter, the reaction mixtures were put in a water bath at 100 °C for 60 minutes to develop colors and the reaction was terminated by incubating the mixtures in ice for 5 minutes. Toluene was added to separate chromophores. The optical density was measured at 520 nm using 1240 UV-Vis Spectrophotometer. Proline content [µmol/g fresh weight (F. WT)] in leaf tissues was calculated from a standard curve made using 0-100 µg L-proline.

### Lipid peroxidation assay

Fresh upper second fully expended leaves (0.3 g) harvested and homogenized in 0.1 % (w/v) trichloroacetic acid and then the homogenates were centrifuged at 10,000 g for 15 minutes at 4 °C. The supernatant was mixed with 0.5 ml of 1.5 ml 0.5% thiobarbituric acid diluted in 20% trichloroacetic acid and the mixture was incubated in water bath at 95 °C for 25 minutes before incubating it on ice for 10 minutes. The absorbance was measured at 532 and 600 nm using UVmini-1240 UV-Vis Spectrophotometer with 1% thiobarbituric acid in 20% trichloroacetic acid as control. The amount of malondialdehyde (µmol/g FW) calculated as a measure of lipid peroxidation, was determined according to Heath and Packer, (1968).

### Estimation of reducing sugar

The amount of reducing sugar in shoots was determined using method describe by Johnson et al (1964). The sugar was extracted from 1.0 g homogenized tissue using 80% ethanol at 95 °C, then centrifuged for 10 min at 14000 rpm. The resulting supernatant was dried for 2 hrs at 80 °C, before dissolving the residue in 10 ml of distilled water and 2.0 ml alkaline copper reagent was added. The mixture was heated in water bath at 100 °C for 10 min, and then cooled to room temperature. Exactly 1.0 ml of Nelson’s reagent was added and the volume was adjusted to 10 ml with double distilled water. Absorbance of the solution was taken at 520 nm. The amount of reducing sugar (mg /g FW) was calculated using a standard curve of glucose.

### Estimation of leaf total protein

Total sample protein was extracted using the acetone-trichloroacetic acid (TCA) precipitation method as described by Damerval et al. (1986). In brief, 500 g of leaf tissue from each treatment was homogenized in 10% TCA in ice and incubated overnight at 4°C. The homogenate was centrifuged at 14,000 rpm for 15 min at 4°C and the pellet was washed with100% acetone to remove any contaminating pigments. To remove phenolic compounds, the pigment-free pellet was first washed with 80% ethanol, ethanol/trichloromethane (3:1 v/v), then ethanol/ethoxyethane (3:1 v/v) and finally with ethoxyethane. The washed pellet was then suspended in a volume of 0.1 N sodium hydroxide for protein estimation. The sample proteins were estimated at 750 nm using bovine serum albumin as standard and expressed as gram per dry weight of tissue.

### Measurements of Na and K and Cl content in plant tissue

Mature leaves from randomly selected finger millet plants were powdered and ashed at 200 °C for 12 hrs. The ashes were dissolved in 5 ml 30% ammonia, and further diluted with deionized water (Cheng et al. 2004). Concentrations of Na^+^ and K^+^ ions were measured using a flame atomic absorption spectrometry. The concentration of chloride ions was determined after aqueous extraction of 1 g of the plant material in 25 ml of distilled water. Concentrations of Cl^-^ ions were determined by titration from the infiltrated solution using silver nitrate in the presence of potassium chromate as described by Eaton et al. (1995).

### Statistical analysis

A completely randomized block design with five replications for each experiment was used and the results represent mean ± standard error. Analysis of variance (ANOVA) was performed using the Minitab statistical computer software version 17 (Minitab Inc., State College, PA, USA) and differences between means were accomplished using the Fisher’s protected LSD test at a confidence level of 95% (p ≤ 0.05). Relationships between the assessed features were performed by Pearson’s correlation. Principal component analysis (PCA) and Cluster analysis (CA) were carry out using the FactoMineR (Factor analysis and data mining with R) package (Husson et al., 2008).

## Results

The present study investigated the changes growth parameters, relative water content, lipid peroxidation level, proline content, reducing sugar and total protein under NaCl induced salinity stress in six finger millet varieties. The parameters analyzed exhibited significant variations among the varieties.

### Effects of salt stress on seed germination

The effect of salinity stress on finger millet seeds germination, evaluated by the percentage of germinated seeds after 17 days, is as shown in Table I. Our results indicate that for all varieties, the germination rate decreased with an increase of the NaCl concentration and varied among the varieties. This decrease in germination rate was most profound at 200 mM and 300mM NaCl concentrations where 0 % germination rate were recorded for all six varieties. In contrast, at moderate stress levels (100 mM NaCl), significant differences in germination profile was observed with GBK043122 having the highest germination rate (46.25%) compared to others whose germination rates ranged from 3.75% to 22.50%. The germination percentage under control conditions was also distinct among the six finger millet varieties and ranged ranging from 90.00% for GBK043137 to 56.25% for GBK043122 (Table 1).

**Table 1.** Effects of NaCl on germination rate of six finger millet varieties

| Variety | Germination rate (%) |  |  |  |
| --- | --- | --- | --- | --- |
|  | 0 mM | 100 mM | 200 mM | 300 mM |
| GBK043137 | 90.0±3.5<sup>a</sup> | 3.8±3.8<sup>c</sup> | 0.0±0.0<sup>a</sup> | 0.0±0.0<sup>a</sup> |
| GBK043128 | 65.0±3.5<sup>bc</sup> | 18.8±2.4<sup>bc</sup> | 0.0±0.0<sup>a</sup> | 0.0±0.0<sup>a</sup> |
| GBK043124 | 80.0±8.4<sup>a</sup> | 37.5±7.8<sup>ab</sup> | 0.0±0.0<sup>a</sup> | 0.0±0.0<sup>a</sup> |
| GBK043122 | 56.3±5.2<sup>c</sup> | 46.3±9.7<sup>a</sup> | 0.0±0.0<sup>a</sup> | 0.0±0.0<sup>a</sup> |
| GBK043094 | 63.8±1.3<sup>bc</sup> | 22.5±11.6<sup>bc</sup> | 0.0±0.0<sup>a</sup> | 0.0±0.0<sup>a</sup> |
| GBK043050 | 76.3±3.7<sup>ab</sup> | 18.85±5.5<sup>bc</sup> | 0.0±0.0<sup>a</sup> | 0.0±0.0<sup>a</sup> |

Values within a column marked with different superscript in each column differ significantly at $p < 0.05$ [Fishers LSD]. Each value represented as mean ± SD are the mean of three replications.

### Growth characteristics in finger millet varieties under salt stress

After phenotypic observation, chlorosis (yellowish color) was observed in all plants under salinity conditions. Leaf chlorosis (yellowish color), leaf scorch, slowed and delayed growth and enlargement of the leaves were distinctly observed in seedlings of all varieties under salinity stress. Plants growing under control conditions exhibited healthy leaves and normal shoot and root developmental stages (Figure 1). The shoot length progressively retarded with increase in NaCl concentration (Table 2). Particularly, the shoot height of GBK043128 population was significantly reduced at the end of under severe salt stress conditions (300 mM NaCl) by about 72.09% while GBK043124 had the least shoot height reduction rate at 63.33% when compared to the control plants (Table 2). Significance variations on the effect of NaCl on shoot length were only observed at 200 mM NaCl concentration. Higher salt concentrations did not record any varietal difference on shoot length (Table 2). Similarly, increasing salinity stress resulted in gradual reductions in plant root lengths in all studied varieties ranging from 20.9% for GBK043137 to 36.1% for GBK043128 compared to their respective controls (Table 3). We also observed significant differences between varieties in root length values across the salt concentrations, signifying that increased salt stress adversely affected root length growth in the varieties at different degrees (Table 3).

**Table 2.** Effect of NaCl on growth of finger millet

| Variety | Seedlings shoot length (cm) under NaCl stress |  |  |  |
| --- | --- | --- | --- | --- |
|  | 0 mM | 100 mM | 200 mM | 300 mM |
| GBK043137 | 3.8±0.4<sup>ab</sup> | 2.5±0.3<sup>a</sup> | 1.6±0.2<sup>ab</sup> | 1.2±0.1<sup>a</sup> |
| GBK043128 | 4.3±0.2<sup>a</sup> | 2.6±0.4<sup>a</sup> | 1.9±0.2<sup>a</sup> | 1.2±0.2<sup>a</sup> |
| GBK043124 | 3.0±0.3<sup>b</sup> | 2.4±0.2<sup>ab</sup> | 1.6±0.3<sup>ab</sup> | 1.1±0.1<sup>a</sup> |
| GBK043122 | 3.2±0.4<sup>b</sup> | 2.5±0.2<sup>a</sup> | 1.2±0.2<sup>bc</sup> | 1.0±0.0<sup>a</sup> |
| GBK043094 | 3.1±0.3<sup>b</sup> | 2.3±0.2<sup>ab</sup> | 1.0±0.0<sup>c</sup> | 1.1±0.1<sup>a</sup> |
| GBK043050 | 3.1±0.2<sup>b</sup> | 1.7±0.1<sup>b</sup> | 1.3±0.2<sup>bc</sup> | 1.1±0.1<sup>a</sup> |

Values within a column marked with different superscript in each column differ significantly at $p < 0.05$ [Fishers LSD]. Each value represented as mean ± SD are the mean of three replications.

**Table 3.** Effect of NaCl on growth root growth

| Variety | Seedlings root length (cm) under NaCl stress |  |  |  |
| --- | --- | --- | --- | --- |
|  | 0 mM | 100 mM | 200 mM | 300 mM |
| GBK043137 | 6.8±0.4<sup>b</sup> | 6.4±0.4<sup>ab</sup> | 5.9±0.2<sup>a</sup> | 5.4±0.2<sup>a</sup> |
| GBK043128 | 8.0±0.4<sup>a</sup> | 7.2±0.4<sup>a</sup> | 5.9±0.3<sup>a</sup> | 5.1±0.3<sup>ab</sup> |
| GBK043124 | 6.9±0.6<sup>b</sup> | 6.3±0.6<sup>ab</sup> | 5.6±0.4<sup>a</sup> | 5.0±0.46<sup>ab</sup> |
| GBK043122 | 6.7±0.2<sup>b</sup> | 6.2±0.5<sup>b</sup> | 5.5±0.3<sup>a</sup> | 4.9±0.2<sup>ab</sup> |
| GBK043094 | 6.8±0.5<sup>b</sup> | 6.2±0.7<sup>b</sup> | 5.9±0.4<sup>a</sup> | 5.4±0.4<sup>a</sup> |
| GBK043050 | 7.1±0.4<sup>b</sup> | 6.4±0.5<sup>ab</sup> | 5.6±0.2<sup>a</sup> | 4.8±0.3<sup>b</sup> |

Values within a column marked with different superscript in each column differ significantly at $p < 0.05$ [Fishers LSD]. Each value represented as mean ± SD are the mean of three replications.

**Fig. 1.**
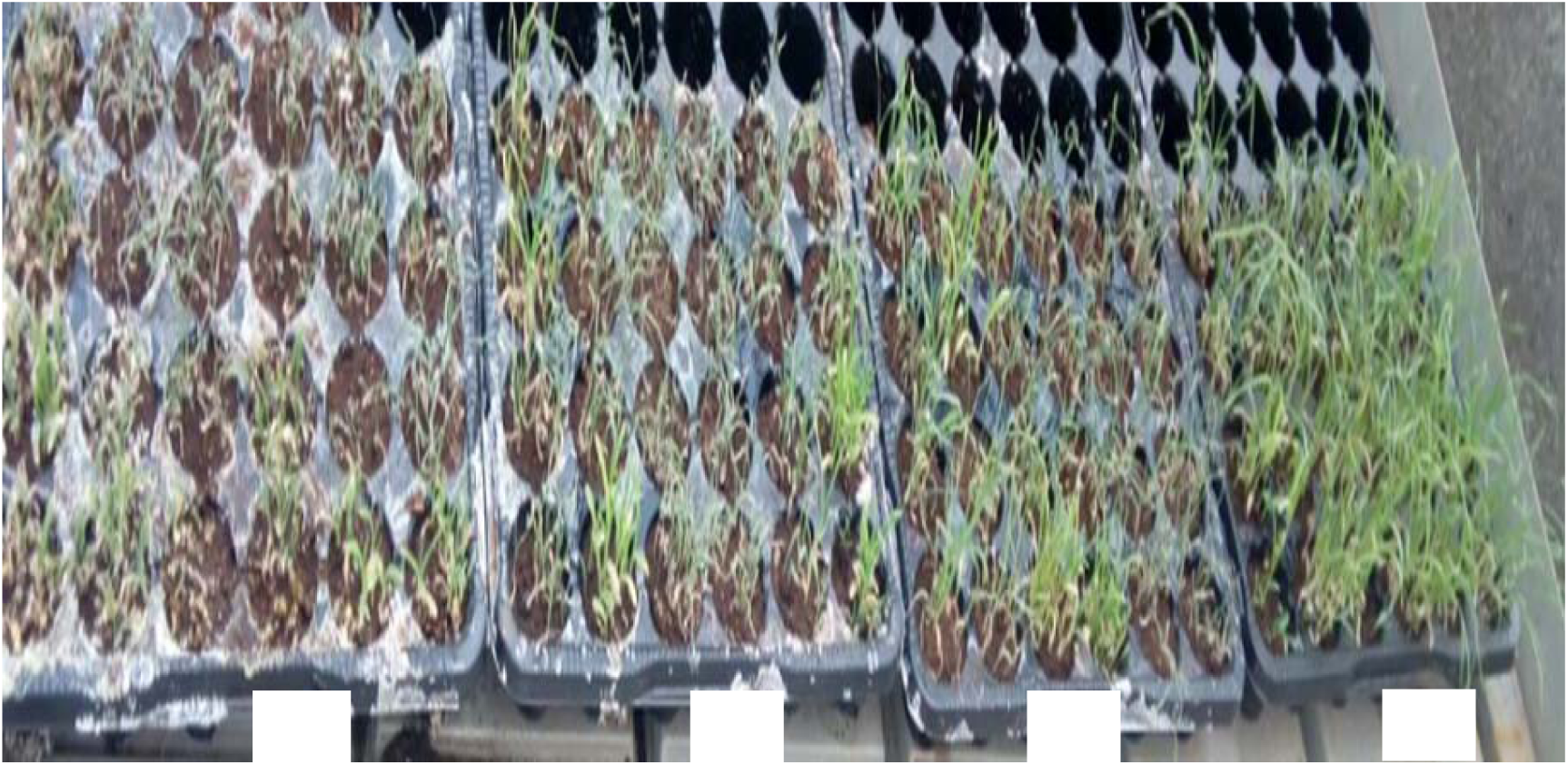
Effect of salinity stress on growth of finger millet. (A) seedling growth on 300 mM NaCl. (B) seedling growth on 200mM NaCl (C) seedling growth on 100 mM NaCl; (D) seedling growth on 0 mM NaCl.

### Relative water content

The changes in leaves RWC along with increase in salinity stress are presented in Table 4. The leaves relative water content of all varieties under control conditions were similar ranging from 79.44 to 87.86%. Exposition to increasing salinity stress progressively reduced water potential of leaves in all varieties compared to their respective control plants leaves and they exhibited variation in their relative water content. Variety GBK043094 tolerated salinity stress better with the least reduction in relative water content under severe salinity stress (300 mM NaCl) compared to the others (Table 4).

**Table 4.** Effect of NaCl on relative water content

| Variety | Seedlings relative water content (%) under NaCl stress |  |  |  |
| --- | --- | --- | --- | --- |
|  | 0 mM | 100 mM | 200 mM | 300 mM |
| GBK043137 | 85.3±4.1<sup>a</sup> | 71.5±4.1<sup>a</sup> | 35.0±3.9<sup>a</sup> | 37.1±3.3<sup>b</sup> |
| GBK043128 | 87.9±5.3<sup>a</sup> | 71.5±4.1<sup>abc</sup> | 35.0±3.9<sup>b</sup> | 26.8±2.3<sup>c</sup> |
| GBK043124 | 84.8±4.9<sup>a</sup> | 67.2±3.4<sup>bc</sup> | 34.2±5.0<sup>b</sup> | 28.2±2.6<sup>c</sup> |
| GBK043122 | 83.0±1.8<sup>a</sup> | 68.0±1.9<sup>bc</sup> | 33.7±3.3<sup>b</sup> | 46.7±9.2<sup>c</sup> |
| GBK043094 | 82.1±6.7<sup>a</sup> | 72.3±3.7<sup>ab</sup> | 51.3±6.1<sup>a</sup> | 46.7±9.2<sup>a</sup> |
| GBK043050 | 79.4±4.6<sup>a</sup> | 65.4±4.8<sup>c</sup> | 33.2±4.5<sup>b</sup> | 27.6±3.7<sup>c</sup> |

Values within a column marked with different superscript in each column differ significantly at $p < 0.05$ [Fishers LSD]. Each value represented as mean ± SD are the mean of three replications.

### Effects of salt stress on chlorophyll content

Analysis of total chlorophyll content demonstrated significant differences in photochemistry among varieties and the salt treatments (Table 5). More specifically, for all the varieties, the addition of NaCl_2_ elicited significant decrease in chlorophyll content compared to the non-saline treatments and inverse relationship between salinity stress and total chlorophyll content in all finger millet varieties was observed. In contrast, plants grown under normal conditions maintained a relatively high levels total chlorophyll content and interestingly, they did not have similar chlorophyll content. Under saline conditions, photosynthetic pigment of varieties GBK043137 and GBK043128 were found to be extremely reduced with reduction percentages of 48.22% and 39.54%, respectively. However, GBK043124 retained a relatively higher chlorophyll content compared to its respective control value, under 300 mM NaCl stress conditions (Table 5). These findings signified that salinity stress may have damaged the photochemical apparatus of the plant leaves.

**Table 5.** Effect of salinity stress on total chlorophyll content of finger millet varieties

| Variety | Seedlings chlorophyll content (mg/g FW) under NaCl stress |  |  |  |
| --- | --- | --- | --- | --- |
|  | 0 mM | 100 mM | 200 mM | 300 mM |
| GBK043137 | 8.4±0.4<sup>a</sup> | 8.1±1.8<sup>a</sup> | 5.0±0.4<sup>a</sup> | 4.4±0.6<sup>a</sup> |
| GBK043128 | 9.1±1.0<sup>b</sup> | 7.5±1.5<sup>b</sup> | 6.4±0.5<sup>b</sup> | 5.5±0.1<sup>b</sup> |
| GBK043124 | 5.9±0.1<sup>c</sup> | 6.8±0.1<sup>c</sup> | 7.3±0.2<sup>c</sup> | 5.5±0.1<sup>c</sup> |
| GBK043122 | 7.3±1.9<sup>d</sup> | 5.9±0.1<sup>d</sup> | 6.1±1.0<sup>d</sup> | 5.8±0.7<sup>d</sup> |
| GBK043094 | 6.2±0.5<sup>e</sup> | 5.0±0.9 | 5.0±0.8<sup>e</sup> | 4.7±1.4<sup>e</sup> |
| GBK043050 | 5.1±1.6<sup>f</sup> | 4.0±0.9<sup>f</sup> | 5.0±0.4<sup>f</sup> | 3.9±0.7<sup>f</sup> |

Values within a column marked with different superscript in each column differ significantly at $p < 0.05$ [Fishers LSD]. Each value represented as mean ± SD are the mean of three replications.

### Proline accumulation and lipid peroxidation assay

Free proline content was estimated in all six finger millet varieties at early seedling growth stage to evaluated their effect under NaCl induced osmotic stress and the data is shown in Table 6. Increasing salt concentrations from 100 to 200 and 300 mM NaCl application remarkably induced increased free proline content in the plants by an average of 1.7-, 2.2- and 3.0-fold change, respectively, relative to the levels in the control plants (Table 6). GBK043094 variety had the significantly highest proline content, followed by GBK043137, GBK043124 and GBK043122 while GBK043128 and GBK043050 had the lowest (Table 6). In unstressed plants, proline concentration was similar. As shown in Table 7, we observed continuous increase in malondialdehyde content in leaves of all varieties tested in response to salinity stress relative to their respective controls and the magnitude of response differed among the varieties. A continuous increase in the level of lipid peroxidation was observed with increasing level of salinity in all the varieties. The malondialdehyde levels (μmol/g FW) was elevated to 20.7%, 31.3% and 51.2% at 100, 200 and 300 mM NaCl, respectively, as compared to unstressed plants (Table 7). Malondialdehyde content was significantly elevated in GBK043050, GBK043122 GBK043124 and GBK043128 under severe salinity stress (300 mM NaCl) treatments signifying higher rates of oxidative damage and lipid peroxidation whereas GBK043094 and GBK043137had lower levels of malondialdehyde at corresponding salinity stress (Table 7).

**Table 6.**
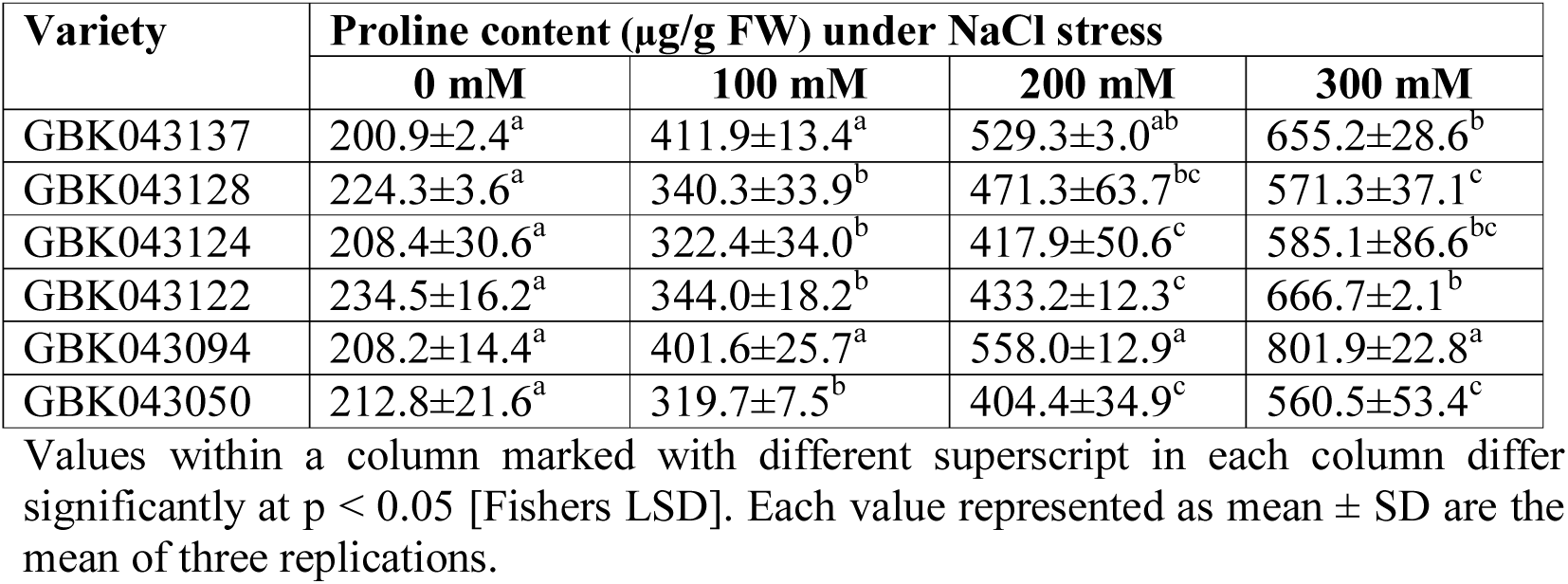
Effect of salinity stress on free proline content of finger millet varieties

| Variety | Proline content (μg/g FW) under NaCl stress | | | |
| --- | --- | --- | --- | --- |
|  | 0 mM | 100 mM | 200 mM | 300 mM |
| GBK043137 | 200.9±2.4<sup>a</sup> | 411.9±13.4<sup>a</sup> | 529.3±3.0<sup>ab</sup> | 655.2±28.6<sup>b</sup> |
| GBK043128 | 224.3±3.6<sup>a</sup> | 340.3±33.9<sup>b</sup> | 471.3±63.7<sup>bc</sup> | 571.3±37.1<sup>c</sup> |
| GBK043124 | 208.4±30.6<sup>a</sup> | 322.4±34.0<sup>b</sup> | 417.9±50.6<sup>c</sup> | 585.1±86.6<sup>bc</sup> |
| GBK043122 | 234.5±16.2<sup>a</sup> | 344.0±18.2<sup>b</sup> | 433.2±12.3<sup>c</sup> | 666.7±2.1<sup>b</sup> |
| GBK043094 | 208.2±14.4<sup>a</sup> | 401.6±25.7<sup>a</sup> | 558.0±12.9<sup>a</sup> | 801.9±22.8<sup>a</sup> |
| GBK043050 | 212.8±21.6<sup>a</sup> | 319.7±7.5<sup>b</sup> | 404.4±34.9<sup>c</sup> | 560.5±53.4<sup>c</sup> |

Values within a column marked with different superscript in each column differ significantly at $p < 0.05$ [Fishers LSD]. Each value represented as mean ± SD are the mean of three replications.

**Table 7.** Effect of salinity stress on free proline content of finger millet varieties

| Variety | Malondialdehyde content (μg/g FW) under NaCl stress | | | |
| --- | --- | --- | --- | --- |
|  | 0 mM | 100 mM | 200 mM | 300 mM |
| GBK043137 | 1.94±0.1<sup>a</sup> | 2.2±0.3<sup>b</sup> | 2.4±0.5<sup>b</sup> | 2.7±0.4<sup>b</sup> |
| GBK043128 | 2.21±0.3<sup>a</sup> | 2.9±0.2<sup>ab</sup> | 2.9±0.4<sup>ab</sup> | 3.6±0.2<sup>a</sup> |
| GBK043124 | 2.67±0.4<sup>a</sup> | 3.3±0.3<sup>a</sup> | 3.4±0.5<sup>a</sup> | 3.7±0.6<sup>a</sup> |
| GBK043122 | 2.22±0.4<sup>a</sup> | 2.8±0.2<sup>ab</sup> | 3.3±0.1<sup>a</sup> | 3.8±0.4<sup>a</sup> |
| GBK043094 | 1.96±0.2<sup>a</sup> | 2.3±0.3<sup>b</sup> | 2.47±0.2<sup>b</sup> | 2.8±0.3<sup>b</sup> |
| GBK043050 | 2.19±0.8<sup>a</sup> | 2.5±0.9<sup>b</sup> | 3.0±0.7<sup>ab</sup> | 3.9±0.5<sup>a</sup> |

Values within a column marked with different superscript in each column differ significantly at $p < 0.05$ [Fishers LSD]. Each value represented as mean ± SD are the mean of three replications.

### Reducing sugars and protein contents under NaCl stress

The impact of salinity treatment triggered substantial elevation in reducing sugar amounts in the stressed plants when compared to control the experiments (Table 8). Increasing salt concentration caused an increase in reducing sugar amounts in the stressed plant shoots and highest accretion of reducing sugar was found in 100 mM NaCl stress followed by 200 mM and 300 mM NaCl treatments. However, varietal differences difference was seen and the increase was remarkably highest in GBK043094, followed by GBK043050, GBK043137and GBK043122 while GBK043128 had the lowest amount (Table 8). Plants under control conditions had the lowest protein content ranging from 1.20 to 2.23 mg/g FW reducing whereas the highest reducing sugar content protein content of 4.47 to 6.45 mg/g FW was found in plants treated with 300 mM NaCl (Table 8). As showed in Table 9, increasing NaCl concentration had a substantial impact on the protein content of finger millet plants and the response was in a dose dependent relationship. A clear varietal difference was observed and significantly higher levels of protein were found in GBK043094, GBK043050 and GBK043122 than the rest, under control and also stress conditions (Table 9).

**Table 8.** Effect of salt stress on reducing sugars on finger millet

| Variety | Reducing sugars content (mg/g FW) under NaCl stress |  |  |  |
| --- | --- | --- | --- | --- |
|  | 0 mM | 100 mM | 200 mM | 300 mM |
| GBK043137 | 1.6±0.4<sup>bc</sup> | 2.1±0.6<sup>ab</sup> | 4.0±0.8<sup>bc</sup> | 4.9±0.9<sup>bc</sup> |
| GBK043128 | 1.2±0.3<sup>c</sup> | 1.6±0.3<sup>b</sup> | 3.7±0.7<sup>bc</sup> | 4.6±0.8<sup>c</sup> |
| GBK043124 | 1.3±0.3<sup>c</sup> | 1.7±0.3<sup>b</sup> | 3.3±0.5<sup>c</sup> | 4.7±0.3<sup>c</sup> |
| GBK043122 | 1.8±0.4<sup>abc</sup> | 2.1±0.4<sup>ab</sup> | 3.7±0.5<sup>bc</sup> | 5.5±0.2<sup>bc</sup> |
| GBK043094 | 2.2±0.3<sup>a</sup> | 2.7±0.2<sup>a</sup> | 5.0±0.0<sup>a</sup> | 6.5±0.5<sup>a</sup> |
| GBK043050 | 2.1±0.4<sup>ab</sup> | 2.5±0.4<sup>a</sup> | 4.4±0.3<sup>ab</sup> | 5.8±0.4<sup>ab</sup> |

Values within a column marked with different superscript in each column differ significantly different at $p < 0.05$ [Fishers LSD]. Each value represented as mean ± SD are the mean of three replications.

**Table 9.** Effect of salt stress on total protein on finger millet

| Variety | Total protein content (mg BSA/g FW) under NaCl stress |  |  |  |
| --- | --- | --- | --- | --- |
|  | 0 mM | 100 mM | 200 mM | 300 mM |
| GBK043137 | 15.2±1.3<sup>b</sup> | 34.4±1.6<sup>b</sup> | 73.8±7.3<sup>c</sup> | 95.7±9.8<sup>b</sup> |
| GBK043128 | 15.3±2.1<sup>b</sup> | 33.9±3.0<sup>b</sup> | 73.1±7.4<sup>c</sup> | 94.7±8.0<sup>b</sup> |
| GBK043124 | 13.2±1.9<sup>b</sup> | 32.1±3.1<sup>b</sup> | 74.7±7.1<sup>bc</sup> | 85.6±4.1<sup>b</sup> |
| GBK043122 | 20.0±2.2<sup>a</sup> | 42.5±5.2<sup>a</sup> | 95.9±4.1<sup>a</sup> | 111.9±7.4<sup>a</sup> |
| GBK043094 | 20.5±3.0<sup>a</sup> | 45.1±5.7<sup>a</sup> | 90.5±9.7<sup>a</sup> | 119.2±6.5<sup>a</sup> |
| GBK043050 | 20.4±1.2<sup>a</sup> | 43.3±3.3<sup>a</sup> | 89.2±11.5<sup>ab</sup> | 117.5±5.4<sup>a</sup> |

Values within a column marked with different superscript in each column differ significantly at $p < 0.05$ [Fishers LSD]. Each value represented as mean ± SD are the mean of three replications.

### Effect of salinity on shoot Na, K and Cl ion composition

The salinity treatments, varieties and the synergy effects were significant for the concentrations of all leaf ions (Fig. 2A, Fig. 2B, Fig. 2C, Supplementary Table 1). As expected, the level of Na^+^ and Cl^-^ in all varieties was higher under salt stress but differed in the degree of the increase. The gradual increase of salinity stress triggered a gradual rise of both ion concentration in finger millet leaves. The average levels of Na^+^ in leaves ranged from 5.37 to 7.82 mg/g DW for plants grown in control conditions and from 12.3 to 96.2 mg/g DW for salinity stressed plants (Fig. 2A). Under 300 mM NaCl stress treatments, the different varieties increased their Na^+^ ion concentration from 6.8- to 13.1-fold when compared to the controls. GBK043124, GBK043137 and GBK043094 displayed statically the minimum increase of Na^+^ under salinity stress (Fig. 2A). On the other hand, the leaf Cl^-^ levels ranged from 2.5 to 5.1 mg/g DW for finger millet plants under control conditions and from 5.0 to 17.8 mg/g DW for plants under salinity stress (Fig. 4). GBK043050 had the lowest concentration of Cl^-^ under untreated and salinity stress treatments. GBK043124 had the least (3.0.5-fold) increase in Cl^-^ ion concentration under salt treatment, while GBK043094 had the largest (4.2-fold) increase ((Fig. 2C). In contrast, salinity stress induced significant reduction of K^+^ concentration in leaves of finger millet plants irrigated with three NaCl doses ((Fig. 3). In comparison to control experiments, potassium ions concentration decreased by about 18.6, 53.3 and 72.6 % in leaves of plants grown under 100, 200 and 300 mM NaCl respectively. GBK043094 upheld the highest concentration of K^+^ and had a 74.0% decline in K^+^ concentration while, GBK043050 had the highest decrease in K content (78.9%) under salinity conditions (Fig. 2B). The lowest potassium ion concentration under salinity was found in GBK043128 followed by GBK043124 (Table 10). The leaf K^+^/Na^+^ ratios differed among the varieties of finger millet studied, ranging from 0.05 in both GBK043094 to 0.02 in GBK043050. Varieties, GBK043094 and GBK043137 presented the greatest K^+^/Na^+^ ratio under salinity stress owing to low concentration of in the leaves (Fig. 2D).

**Fig. 2.**
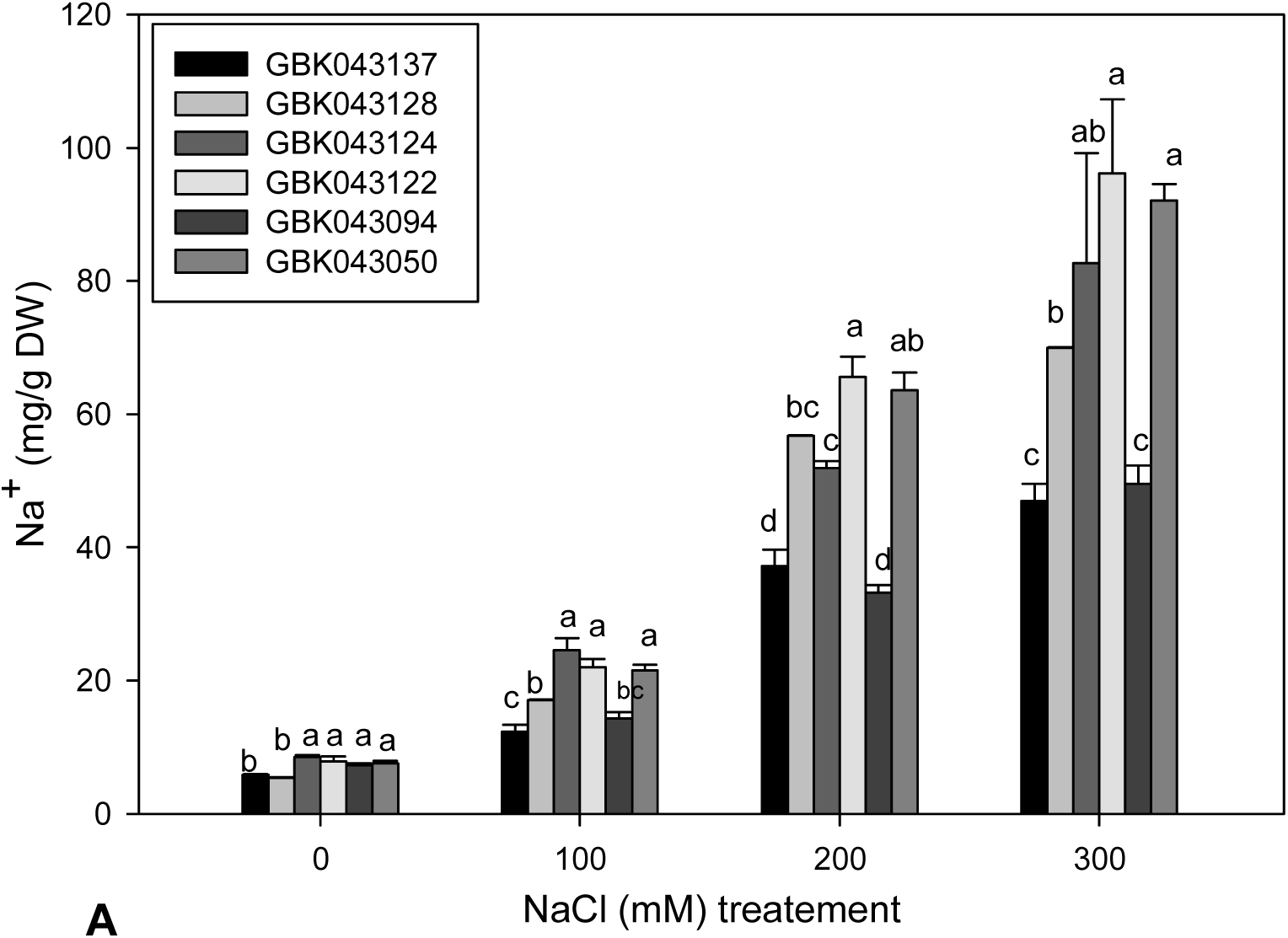
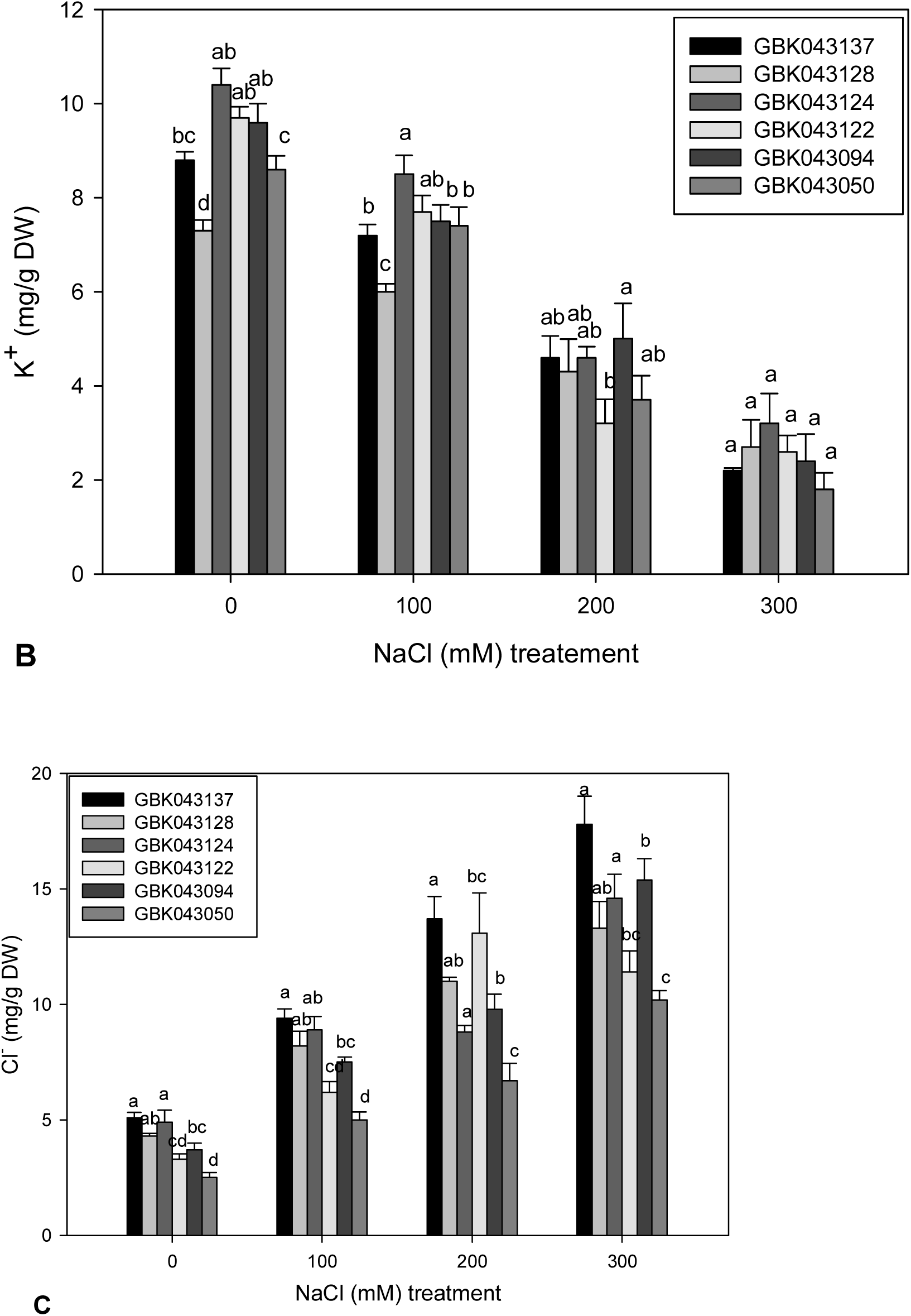
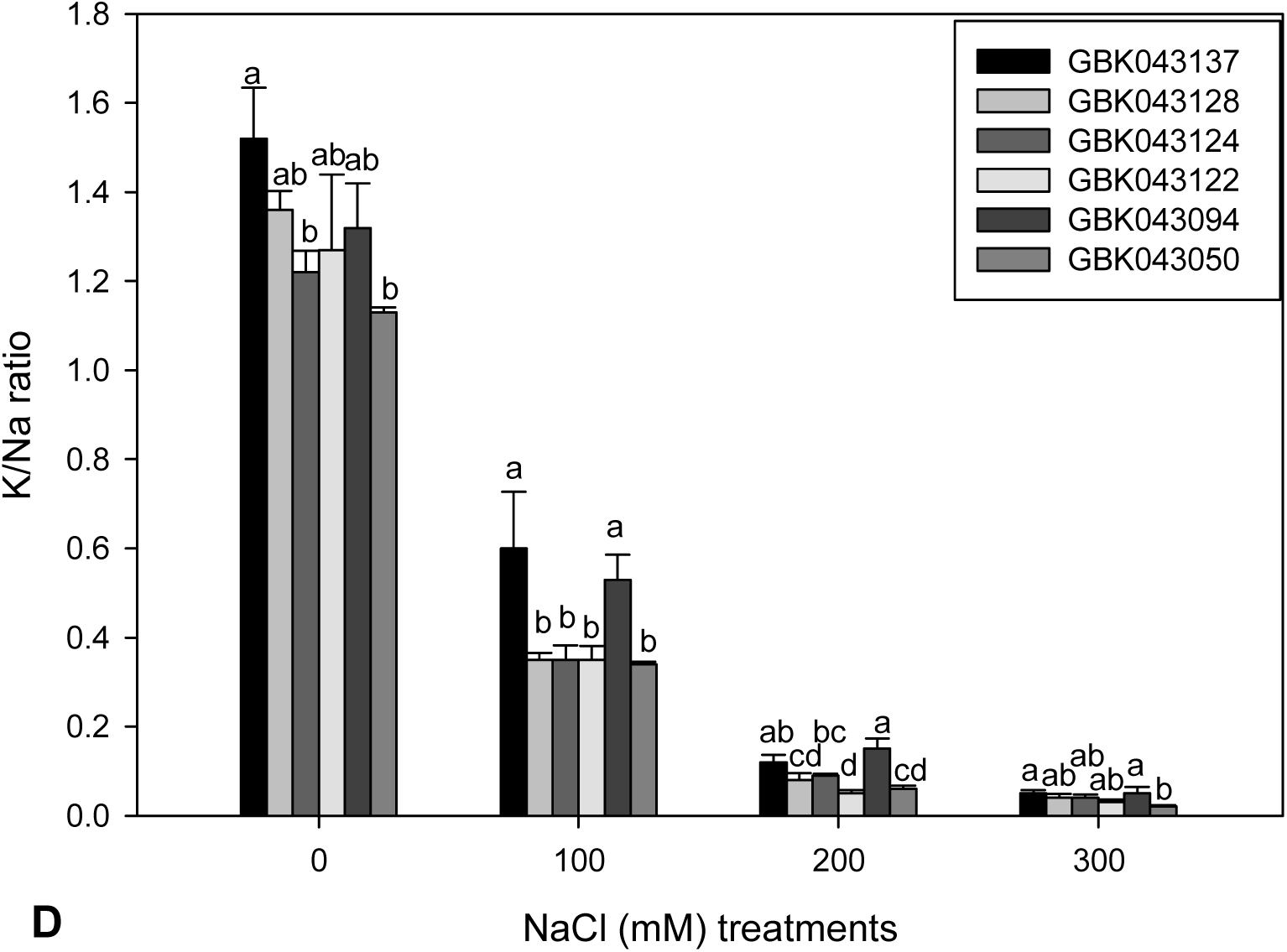
Effect of salinity stress on ion concentration of finger millet under salinity stress. **A:** Na^+^ concentration, **B** K^+^ concentration, **C Cl**^**-**^ **concentration, D** K^+^/Na^+^ **ratio**

**Fig. 2.**
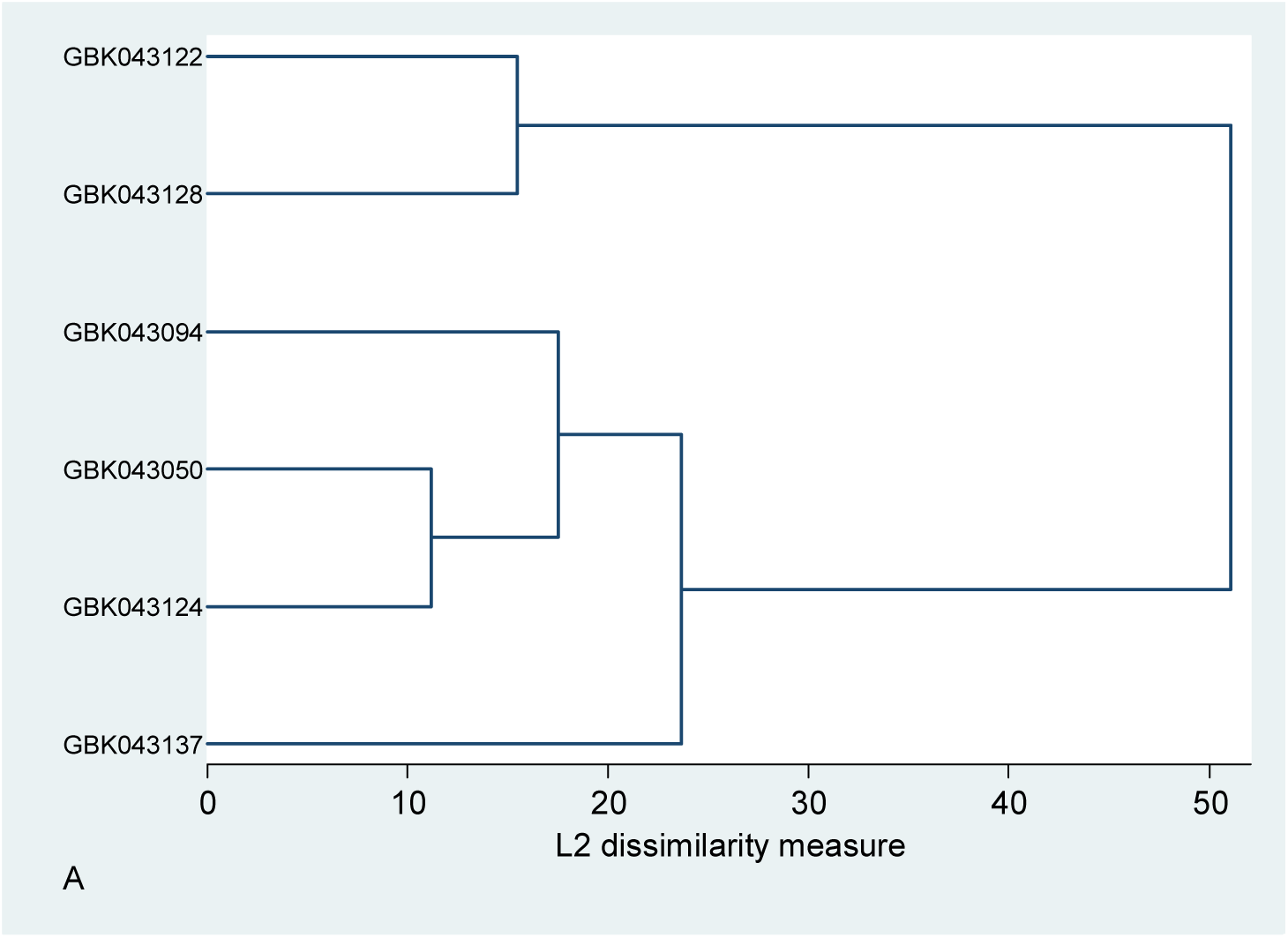
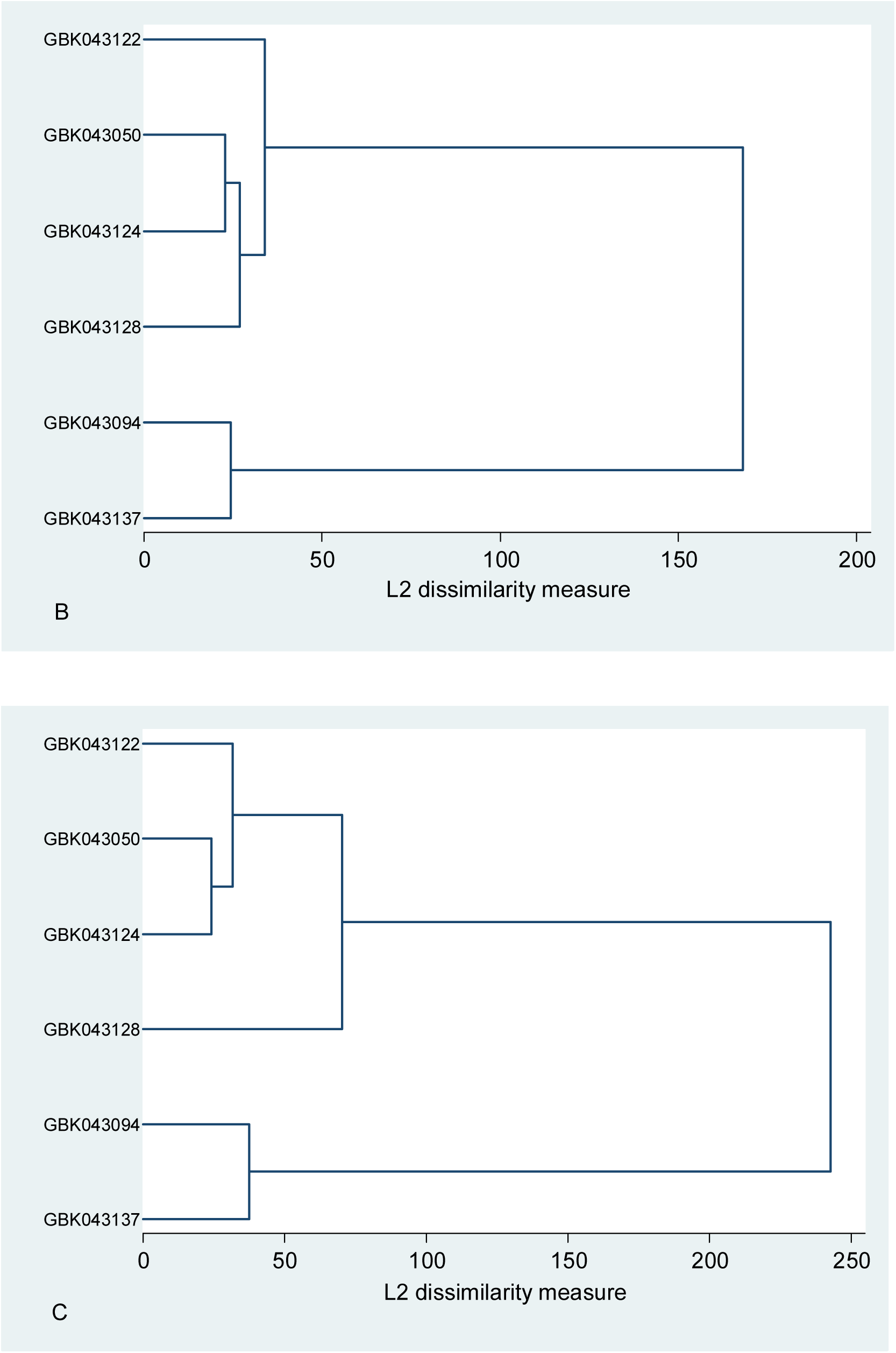
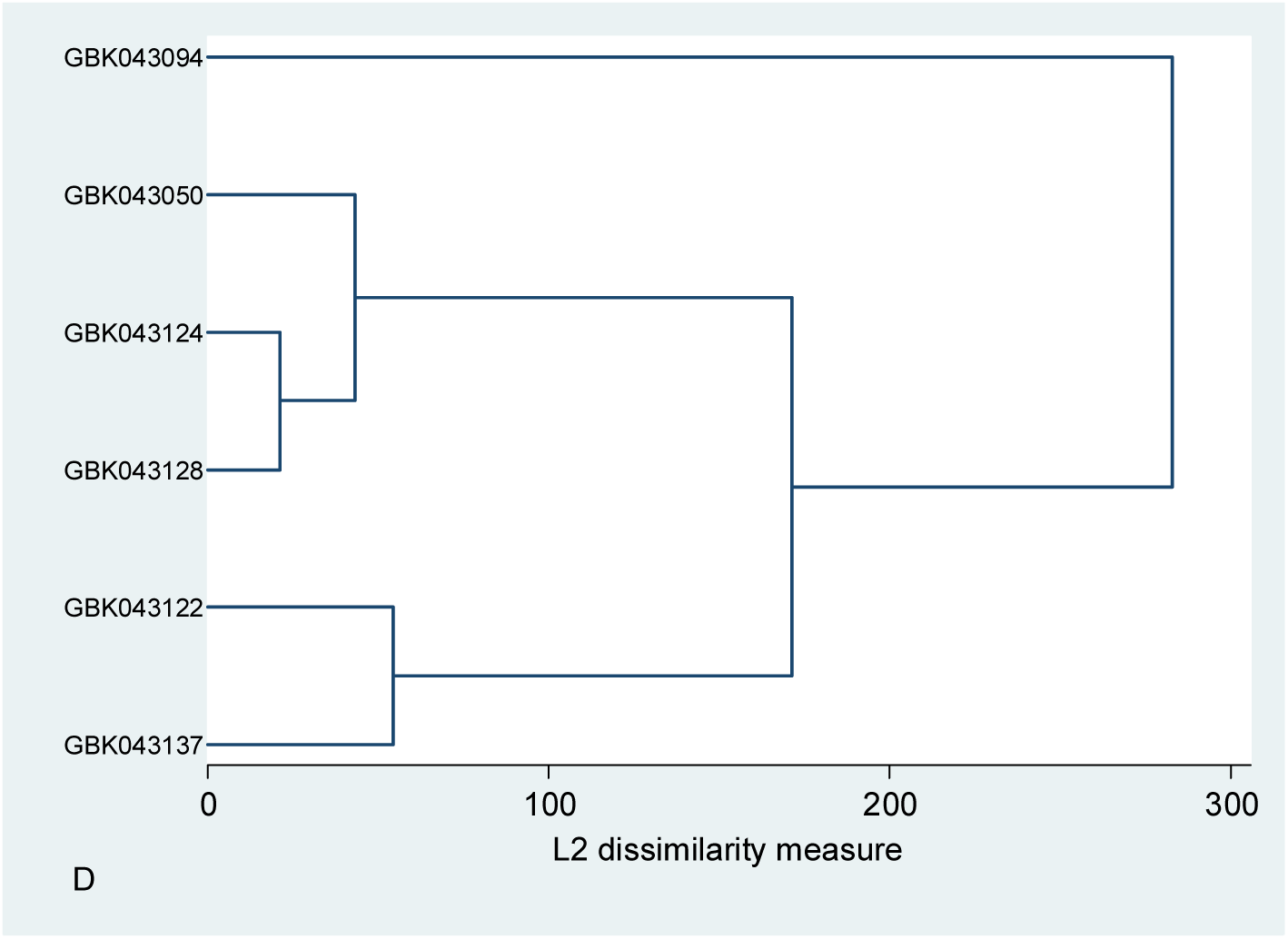
Dendrogram of the studied finger millet varieties, obtained by cluster analysis based on their physiological and biochemical characteristics under salinity stress.

### Cluster analysis

Cluster analysis using average linkage method of clustering was done to classify the varieties into homogenous groups using the physiological and biochemical traits of control and salinity stress treatments. Clustering grouped the six finger varieties into two major clusters based to their potential characteristics under control and salinity stress conditions, respectively (Fig. 2A, B, C and D). Varieties grouped into specific classes indicate the presence of greater diversity among finger millets under different salinity stresses, with varieties GBK043137 GBK043094 showing greater tolerance to salinity stress.

## Discussion

Plants tolerance to salinity stress is a complex trait which is ascribed to a plethora of related morphological, physiological and biochemical adaptive responses and operate synergistically to lessen cell hyperosmolarity and the ensuing ion disequilibrium (Parihar et al., 2015). In this regard, screening and selection finger millet varieties tolerant to salinity stress is essential in order to understand their adaptations under saline soils and for successful production of finger millet in salinity prone areas. In this study, six finger millet varieties from different agroecological zones in Kenya were subjected to different levels of salinity stress, and our findings show tremendous variabilities occur within the tested parameters.

Seed germination and seedling emergence are fundamental biological processes in plant growth and development cycles, and therefore excellent seed germination and emergence are important for attainment of high yields and increasing concentrations of salt adversely affects germination process (Laghmouchi et al., 2017; Anuradha et al., 2001). In the present study, the germination percentage was delayed or constrained under salinity stress compared to control growth conditions. The observed decrease in germination rate under the salinity stress could be attributed to salt toxicity and changes in cellular osmotic potential. We found out that Under higher high hypertonic potential, the reduction in the germination rate was less for the salinity-tolerant variety (GBK043094), compared with most salt sensitive variety (GBK043050). Our finding was in accordance to previous work in lettuce (Ahmed et al., 2019), alfalfa (Sandhu et al., 2017) and wheat (Tounsi et al., 2017) under saline conditions. In addition, a high degree of shoot growth depression in seedlings grown under salinity stress was clearly noticeable, more in the salt-sensitive varieties, which displayed reduced leaf area, leaf chlorosis, leaf burns and plant death, symptoms associated with plant toxicity. Slower growth of both shoots and roots is an adaptive characteristic for plant survival under salinity conditions because this permits the plants to commit numerous resources to mitigate the stress (Soares et al., 2018). Retarded shoot growth under salinity stress could be ascribed to the reduction in osmotic potential due to extra concentration of sodium and chloride ions in the shoot and root zone resulting to a nutritional imbalance and also due to the deviation of energy destined for growth and development to exclude sodium ions cellular absorption and biosynthesis of solutes for preservation cell turgor during hypertonic saline conditions. The observed reduction of leaf area under salinity treatments compared to control plants also suggests that salinity stress may affect plant growth through reduction in leaf area. Previous works have disclosed that salt tolerant plants displays less growth retardation and have relatively higher growth rate compared to sensitive ones under salinity stress (Carillo et al., 2019; Hussain et al., 2018; Sarabi et al., 2017;). Consequently, our findings suggest that GBK043128 and GBK043137 have a better capacity to sustain growth and development under salt treatments compared to other finger millet varieties studied, to sustain growth and production under salinity conditions (Table 2). Further, roots are often reported to play a key role in the salt tolerance of plants as they represent the first organs that control the uptake and translocation of nutrients and salts throughout the plant. Because of their direct exposure to saline environment, root growth is also vulnerable to salt stress although the extend is less than that of the shoots (Munns and Tester, 2008). The inhibition of root growth in plants adversely affects the survival and productivity of the plants and therefore, root growth under saline conditions may serve as good indicator in the first steps of screening for salinity tolerance programs. The growth of roots varies widely due to soil conditions because the status of all nutrient in plants is maintained from the soil with the help of roots. Root growth rate may be severely affected by saline soils and reduction may even be recorded in salt-tolerant plants. In agreement with previously published studies on the effects of salinity on root elongation (Cirillo et al., 2019; Dugasa et al., 2019), salinity treatments were found to cause stunted root growth. The growth-promoting effect under salinity stress could be due to an increase in the osmotic potential of the cells in the elongation zone coupled with enhanced cell division. The absorbed ions at this point could be quickly compartmentalized into the vacuoles without getting to the maximum capacity, thereby increasing the turgor within the cells and stimulating cell elongation. We also observed that the effects of salinity stress on root grow was much less compared to that of the shoot. This feature could be explained by the fact that roots are less affected by salt salinity due to transport of ions to other plant organs and hence the stressed roots to maintain osmotic balance.

Accumulation of ions in plant tissues is regularly used to evaluate the capability of a plant to resist salt stress and salinity is known to cause fluctuations of macronutrients. The concentration of sodium, potassium and calcium ions and the K/Na ratio are vital features that usually used for screening of salt tolerant plants (Sarabi et al., 2017). We used leaf tissues because they are more sensitive to salt and start displaying toxicity much earlier compared to other plant organs (Munns and Tester, 2008). Our study revealed that the NaCl treatments increased the finger millet leaf Na^+^ and Cl^-^ concentrations. Contrary, salinity treatments caused decease of K^+^ in all varieties, probably due to membranes depolarization and loss of Ca^+^ ion due to the displacement by Na^+^ ions. It has also been established that Na^+^ and K^+^ have similar cellular effects despite the fact Na^+^ inhibits K^+^ absorption through binding and obstructing to its transport system (Flowers and Yeo, 1986). It has been established that Na^+^ and K^+^ have similar cellular effects despite the fact Na^+^ inhibits K^+^ absorption through binding and obstructing to its transport system (Flowers and Yeo (1986). Many studies have reported that plants growing under high NaCl concentrations have low ratios of K^+^/Na^+^ ratio caused by deficiency of intracellular K^+^ (Dugasa et al., 2018; Cirillo et al., 2018; Sandhu et al., 2017; Sarabi et al., 2017). The same phenomenon was also observed in this study, where increment of NaCl concentration decreased the leaf K^+^/Na^+^ ratios. In our study, we observed a clear association between K^+^/Na^+^ ratio and salinity tolerance and varieties, GBK043137 and GBK043094 showed the highest K^+^/Na^+^ ratios under both control and NaCl treatments, however, these varieties were placed at the highest ranking for salinity tolerance index. Usually, cellular influx of Cl^-^ ions influx require energy in a reaction mechanism catalysed by a Cl^−^/2H^+^ coupled antiporters and symporters and it is typically taken up freely with water uptake, and is therefore accumulated in leaf organs depending on the transpiration rate (Munns and Tester, 2008). Like Na^+^, Cl^−^ ions may also be sequestered in cell vacuoles. In our study, the concentration of Cl^-^ in leaves was higher than that of Na^+^ and this may be justified by the partial control of Na^+,^ at roots. Comparable results were also exhibited by melon (Sarabi et al., 2017) and cucumber (Colla et al., 2012).

Several studies suggest chlorophyll content as a biochemical marker of salt tolerance in plants (Ishikawa Shabala, 2019; Taïbi et al., 2016; Sairam et al., 2005). It is known that salt tolerant plants show increased or unchanged chlorophyll levels under salinity conditions whereas chlorophyll contents decreased in salt-sensitive plants (Stepien and Johonson, 2009; Ashraf and Harris, 2013). In general, decrease of chlorophyll content under salt stress is considered to be a result of slow synthesis or fast breakdown of the pigments in cells (Ashraf, 2003). The decrease in total chlorophyll content may also be observed due to ion accumulation and functional disorders observed during stoma opening and closing under salinity stress (Nawaz et al., 2010). Another reason for the decrease of chlorophyll content under salt conditions is stated to be the rapid maturing of leaves (Yeo et al., 1991). In our study statistically significant decrease in total chlorophyll content was observed with increasing salt concentration. Similar results were reported by Ashraf and Yousafali (1998) and Ali et al., (2004) and showed that the total chlorophyll content of rice leaves was generally reduced under high salinity. While the other varieties recorded a decrease over the control plants, variety GBK043094 recorded unchanged total chlorophyll content with increase in stress (Table4). These results showed that the reduction in chlorophyll content was variety specific and some varieties showed comparatively lesser quantum of negative variation in chlorophyll content thus indicating their potential to grow and perform moderately well even under higher levels of salt stress. High salt concentration induced reduction of total chlorophyll content indicates that salt stress induces chlorophyll degradation and destruction of chloroplast structures.

All plants employ complex biochemical defensive mechanisms against oxidative injury of free radicals and ROS during abiotic stresses. Among these defence systems is the aggregation of compatible solutes such as proline, an osmoprotectant that preserves membrane integrity and mitigates oxidative burst in plant challenged by salt stress Ahmed et al. 2013, Rao et al., 2013). In addition, proline exists in all plant organs, accumulating in greater proportions compared to other amino acids in salinity stressed plants (Banu et al., 2009). Although the beneficial outcome of proline overproduction in plants during salinity stress have been explicated, the definite roles of proline accretion are still obscure (Banu et al., 2009; Verbruggen and Hermans, 2008). Our study reported an increased concentration of free proline content in all six finger millet varieties with GBK043094 and GBK043137 displaying higher free proline amounts at all salinity treatments suggesting that they are comparatively more tolerant to salinity stress than the rest and which may be related to their competitive ability under saline stress against oxidative stress. Based on these results, it is worth noting that increased concentration of free proline content in finger millet plants subjected to salinity treatments corresponded to improved salinity tolerance. Degradation of polyunsaturated fatty acids in plants yields malondialdehyde (MDA) a biomarker for determining the degree of lipid peroxidation and cellular membrane (Yang et al., 2018). Results from our study reveals that MDA the content in stressed plants raised with increasing stress levels corroborate with those exhibited in other plant species like *Lycium ruthenicum* (Li et al., 2019), wheat (Dugasa et al., 2018) and *Cucumis melo* L. (Sarabi et al., 2017). Our results indicated that some varieties finger millet may tolerate saline environments than others depending on the severity of the stress, by lowering the rate of lipid peroxidation and the cell membrane damage and therefore have an efficient and effective antioxidant defence mechanism. Moreover, the strong negative correlation witnessed between MDA and shoot height (r= – 0.6872, Supplementary Material 1), and root length (r= –7555, Supplementary Material 1) affirms that the NaCl stress triggered lipid peroxidation is one of the reasons for the observed stunted shoot and root growth in finger millet plants.

Contrary to other osmolytes such as proline and MDA which are present at very low amounts except when their biosynthesis is triggered by stress, compatible solutes such as reducing sugars are elements of metabolism with different cell functional roles, such as precursors of other metabolites, signalling molecules and major source of energy. Their levels are highly controlled by various systems to ensure cellular homeostasis. Reducing sugars therefore play a crucial role in plant cells osmotic adjustment during salinity stress. The higher reducing sugar levels measured plants with high salinity tolerant index clearly shows that the sugar contributes to osmotic adjustment during salt treatments thus cushioning the plants against the toxic effects of NaCl. These results are substantiated by a remarkable increase in sugar amounts in salt tolerant genotypes in pigeon pea (Awana et al., 2019), *Juncus* sp (Hassan et al., 2016) and wheat (Kerepesi and Galiba, 2000). Likewise, accumulation of protein compounds has essential part in physiological responses of plant to salinity stress. Increased production of proteins and other nitrogen containing compound may induce the biosynthesis of osmotically active organic compounds including proteins with osmoprotective capacities, thereby conferring salinity resistance (Ashraf and Harris, 2004). Generally, plants exposed to NaCl stress have comparatively reduced protein levels which often results to loss of cellular turgor. Just like in our case, reduction in the content of soluble protein was observed in maize plants subjected to salinity treatments (von Alvensleben et al., 2013).

Lastly, it is imperative to note that the results of this study were conducted in a laboratory set-up (artificial conditions), which may not mirror their complex natural. However, the findings give suggestive index salinity tolerance to the studied finger millet varieties.

## Conclusions

This study gives a deep analysis of the effect of a NaCl stress treatments on the physiological and biochemical parameters six finger millet varieties. In conclusion, our results demonstrated salinity responses on the evaluated features with significant varietal differences among the plants studied and supported by observations made. From the responses of GBK043094 and GBK043137 varieties, we hypothesised that these varieties are promising genetic resources with comparative high tolerance to salinity and hence they may be utilised for further assessment for breeding programs of the crop towards enhanced salinity tolerance. Our findings give suggestive salinity tolerance index to the studied finger millet varieties and should be confirmed for in a wide range a wide range of environmental conditions and other salt types. Lastly, it is imperative to note that the results of this study were conducted in a laboratory set-up (artificial conditions), which may not mirror their complex natural environments under which the crop is grown.

## Acknowledgements

This project was funded by The World Academy of Sciences grant (Ref. No. 17-357 RG/BIO/AF/AC_I – FR3240297745) through the generous contribution of the Swedish International Development Cooperation Agency. The authors thank the Kenyatta University and Pwani University for providing laboratory facilities. The authors gratefully acknowledge Kenya Agricultural and Livestock Research Organization, Gene Bank, for providing the finger millet seeds.

## Author contribution

Asunta Mukami, Alex Ngetich, Wilton Mbinda designed and performed the experiments performed data analyses, Asunta Mukami, Wilton Mbinda wrote the draft manuscript., Easter Syombua and Richard Oduor revised and corrected the draft manuscript, Wilton Mbinda conceptualized the idea and design, supervised the work and made critical review of the article.

